# Black women in medical education publishing: Bibliometric and testimonio accounts using intersectionality methodology

**DOI:** 10.1101/2023.08.31.555784

**Authors:** Witzard Seide, Lauren A. Maggio, Anthony R. Artino, Todd Leroux, Abigail Konopasky

## Abstract

**Background:** Black women in academic medicine experience racial and gender discrimination, all while being tasked with improving a flawed system. Representation of Black women in medicine remains low, yet they bear the burden of fostering diversity and mentoring trainees, exacerbating their minority tax and emotional labor, and negatively impacting career progression.

**Objective:** To complement qualitative accounts of Black women authors in the medical education literature with a quantitative account of their representation.

**Design:** An intersectional methodology employing bibliometric analysis and testimonio reflection.

**Subjects:** US-based authors of journal articles published in medical education journals between 2000 and 2020.

**Main measures:** Author race was determined using a probability-based algorithm incorporating US Census data, and author gender was ascribed using Social Security Administration records. We conducted two negative binomial generalized linear models by first and last author publications. Metadata for each article was retrieved from Web of Science and PubMed to include author names, country of institutional affiliation, and Medical Subject Headings (MeSH). Results were contextualized via the “testimonio” account of a Black woman author.

**Key Results:** Of 21,945 unique authors, Black women (and other racially minoritized groups) published far fewer first and last authors papers than white women and men. In addition, major MeSH terms used by Black women authors reveal little overlap of highly ranked medical education topics. The testimonio further narrated struggles with belonging and racial identity.

**Conclusion:** Black women are underrepresented in medical education publishing and feel a lack of belonging. We believe that dismantling oppressive structures in the publishing ecosystem and the field is imperative for achieving equity. Additionally, further experiential accounts are needed to contextualize this quantitative account and understand oppression in medical education publishing.

## Introduction

To be a Black woman in academic medicine is to experience racial and gender discrimination and otherness, “drowning in the same system we are burdened to improve.”^1(p1)^ Black women are called upon to mend this broken system, by recruiting and mentoring trainees of color and leading diversity efforts; this minority tax burden,^1^ in addition to requiring that Black women perpetually navigate the emotional labor to “anticipate and deflect harm” ^2(p329)^ is heavy, since Black women comprise only 1.5% of medical school faculty.^3^

Some Black women scholars are starting to speak out, specifically about their experiences of discrimination in academic *publishing*, a critical component of academic career progression. ^4–6^ Their stories argue for the *experiential significance* of this discrimination, but as a field steeped in quantitative methodologies and a postpositivist desire for generalizability, *statistical significance* is an important aspect of arguing for change.^7^ Current literature has used quantitative methodologies to explore author gender,^8,9^ but it has not done the same for race/racism (following Gilborn and colleagues, we are “ambivalent” about race: it has very real effects but they come through mechanisms of rac*ism*; we denote this by using these two words together).^10^ Here we add to Black women authors’ *experiential* accounts in medical education publishing with a *statistical* account. We do so to support and center these experiential accounts, not supplant them.

The literature contains both experiential and statistical accounts of discrimination against Black women in medicine writ large. For instance, a gastroenterologist in academic medicine shared experiences of racism and misogyny, noting the pervasiveness of this “Intersectional discrimination.” ^1(p1)^ In another recent commentary, authors discussed how as Black women in academic medicine, they have to, “work twice as hard to get half as far” ^11(p1398)^ without any safety net. They characterize this as the “Goldilocks dilemma: [Black women] are either insufficient and unsuitable or boastful and overdone–never just right.” ^11(p1399)^ The literature also provides a robust statistical account: Black women comprise just 28% of medical school assistant professors, 1.8% of associate, and a mere .76% of full professors.^3^

Unfortunately, we have less robust accounts in regards to *publishing*. Two recently published experiential accounts point to Black women’s vulnerability in the peer-review process in academic medicine: comments from reviewers may express bias when authors reveal their identity,^5^ and this can lead Black women authors to feel like their story is “not good enough.”^4(p145)^ Statements from editorial leaders in medical education support these accounts, noting that bias is baked into the publishing process.^12,13^ Yet there are no intersectional quantitative accounts to stand alongside those experiences. Studies focusing *separately on race* in publishing have found that: Black authors publish more articles on disparities, whereas white authors publish across all topics^14^; this has career implications as different topics receive more or less citations.^15^ Finally, the citation impact of Black authors lags behind others.^16^ Meanwhile quantitative accounts focused *separately on gender* have found that: women tend to publish qualitative articles on topics that are people-related (e.g., caring for others);^17^ men tend to publish quantitative articles on topics related to things (e.g., power; politics; business)^18–20^ Additionally, there are gender-based inequities in citation patterns (i.e., white men being more heavily cited).^20^ Finally, in one quantitative, intersectional study of inequalities in scientific publishing the authors found that Black women were less cited than white men in the fields of physics, mathematics, and engineering and less cited than white men and women in lower-cited fields.^14^ We used these methods in the present study to offer a statistical account of Black women in *medical education* publishing to complement previous experiential accounts.

### Approach

Most accounts of publication inequities examine racism and sexism separately. This misses the complex “intersectionality” first noted by Kimberlé Crenshaw, who noted the reciprocal and multiplicative functioning of racism for Black women, “shaping structural, political, and representational aspects of violence.”^21(p1244)^ In quantitative accounts, narrating this complexity is difficult for several reasons. First, statistical methods rely on separate variables and robust numbers and, as noted above, there are few Black women in medical education publishing. Second, the history and practice of quantitative methods is intertwined with racism: two of its founders, Galton and Pearson, developed statistical methods largely for eugenics research.^22^ Third, assumptions of objectivity and generalizability work against explaining the contextualized and lived experiences of Black women.^10,23^

Nonetheless, quantitative approaches are valued in medical education, and they offer a different perspective that we believe can complement, rather than supplant, experiential ones. Moreover, publications are central sources of advancement and prestige.^4–6^ Therefore, understanding their (in)equitable distribution is critical to making change. We center Black women’s intersectional experiences using Intersectionality Methodology (IM),^24^ developed by Black women scholars through analysis of intersectional studies of Black women. We also draw on Zuberi and Bonilla-Silva’s *White Logic, White Methods*^25^ and the ensuing “QuantCrit” movement.^23^

### Study Purpose

The purpose of this study is to offer a statistical account of Black women in medical education publishing to complement existing experiential accounts and, critically to IM, to contextualize this account through the experiences of a Black woman author in medical education.

## Methods

IM emphasizes (1) power, (2) reflexivity (especially for researchers who are not Black women), (3) centering Black women through statistical methods, and (4) “presenting Black women in the fullness of their humanity”.^24(p778)^ We present our methods below using this IM framework.

### Power

IM asks researchers to attend to power relations, addressing how they are at work in the research process. Medical education is dominated by whiteness, which acts as a gatekeeper to keep Black women and others out.^26^ This is evidenced by Black women’s accounts of oppression and exclusion in medical education^1,11,27,28^ and publishing.^4,5,29^ This power is also evident in the field’s reliance on biomedical inquiry, with its sole focus on the body to the exclusion of political and social structures^30^. For instance, the term “underrepresented in medicine” both denotes racially minoritized physicians and trainees like Black women as ”other” and homogenizes their experiences.^31^ The statistical account below is inextricably tied to this context of power.

To mitigate this power, we present some of our limitations here rather than at the end of the paper, to highlight the inequity of our data sources. First, bibliometric tools like the Web of Science (WoS) are created predominantly by male peer reviewers^32^ and white editorial boards.^33^ Second, the US Census was created to protect white individuals and still retains limitations of that history.^34^ Finally, the Social Security Administration (SSA) has a racist history, with arguments that it disproportionately benefits whites.^35^

### Reflexivity

Our research team identities also reflect power inequities. Indeed, the first author, a researcher with an MD who is a uniformed services officer, is the only Black woman (or Black) member of our team and, as such, the only one with intersectional experiences. The four other authors include two white women and two white men, all PhD-level researchers who have benefited from racism. This imbalance reflects the structural inequities of the medical education research field in which we met, but also reflects how the white academics on the team have not necessarily seen or valued their Black women colleagues appropriately. While we used IM and reflexivity to continually *re*center the first author and the first author herself has intentionally ensured she is empowered to speak up and that her voice is heard throughout this manuscript, we know that the results below are limited by the racial lenses of the four white team members.

### Centering Black Women in a Quantitative Bibliometric Analysis

We conducted a bibliometric analysis^36^ of US-based journal article authors focused on their predicted gender and race/racism. This project is a sub-analysis of a larger project that broadly investigated author characteristics globally, not attempting to identify race/racism.^9^ In the current study, we utilized a subset of the data specific to US-based authors. The data set drew from the Medical Education Journal List (MEJ-24), described as a “seed set of journals” representing the field of medical education.^37^ The MEJ-24 was derived using co-citation, an evidence-based approach for field delineation (while also created largely by those in positions of publishing power).^38–40^ All data assembly, cleaning, and statistical modeling were performed using R version 4.2.1.

While we seek to support experiential accounts of Black women with a statistical account, the bibliometric and statistical methods we use are themselves oppressive, steeped in the white logic of the academy.^10,25^ We draw four tools from QuantCrit to mitigate this: (a) whenever possible (i.e., when it will not be misrepresentative), we foreground intersectional identities^7^ (b) we always refer to racism alongside race, (c) we contextualize our work around power and reflexivity (see sections above),^7,23^ and (d) our first author offers a *testimonio* – “purposeful storytelling grounded in praxis utilized to expose and disrupt histories that are otherwise subsumed”^7(p255)^ --to center lived experience.

### Data

On August 27, 2021 we used WoS to retrieve metadata for articles published in 22 of the MEJ-24 journals between 2000-2020 (as the *Journal of Graduate Medical Education* and the *Canadian Medical Education Journal* are not indexed in WoS). On the same day, we downloaded metadata for all MEJ-24 journals using the Crossref REST API. Metadata included author first and last names, journal, publication year, title, and abstract. For all journals we downloaded the associated Medical Subject Headings (MeSH), which we use as a proxy for article topic (recognizing the limitations from its origination in the oppressive structure of publishing). From this data set we identified and extracted all authors affiliated with institutions in the United States (US). We necessarily focused on US-based authors because our approach utilizes US Census data to provide the basis for making estimations on author race/racism. Two separate algorithms were utilized to provide estimates on race/racism and gender for the authors captured in these journals. In keeping with IM, we present and interpret our findings as an *intersection* of these algorithms.

### Race/Racism Algorithm

We adapted a race/racism algorithm originally developed by Kozlowski et al.,^14^ which utilized family names (surnames) and given names, with data on frequency counts by the US Census Bureau, to assign probabilities of race/racism based upon these names. The algorithm collapsed categories of Asian*, Black*, Hispanic*, white, and other* (these categories are not necessarily reflective of the lived identity of those to whom they refer, but we use the Census terms in order not to misrepresent the data set; we henceforth mark them with an asterisk to note their potential to oppress). We collapsed some race/racism groupings from the 2010 Census to increase power for estimating effect sizes. Of note, two categories were pooled (“Other*”) when performing summary statistics and statistical models due to low sample sizes (Non-Hispanic American Indian* and Alaska Native Alone* and Non-Hispanic Two or More Races*).

A key element of this algorithm is that, rather than utilizing a race/racism assignment of authors to the most prominent racial group, each author was given an associated *probability* to each race/racism classification (i.e., the sum of all race/racism categories, by author, sums to 100%). Previous research demonstrated^14^ that assigning race/racism based upon the most prominent racial group (or an overall attribution), often underestimates Black* authors and overestimates white authors. We used this algorithm with assigned probabilities across all racial categories to examine intersectionality at the aggregate level, not the individual author level. In cases of missing data, if the Census data did not capture the family and given name, mean imputation was used for the author’s racial probability based upon the distribution for the entire analytic sample.

### Gender Algorithm

While gender is neither fixed nor binary, it is a powerful social construct, so we used the categories “women” and “men” here to explore sexism. To classify authors by gender, given name data covering years 1880-2021 from the SSA^41^ were used to assign gender categories to authors based upon the frequency of the name from birth records. These data captured the top 1,000 names for each year, and each name is associated with a binary gender (women or men). Unlike the above racial attribution methodology, here, authors were classified as women or men according to which name in the SSA data had the greatest frequency attributed to the specific gender rather than assigned a probability.

### Statistical Analysis

Two negative binomial generalized linear models were used to estimate the average effect of epistemic oppression on Black* women, from 2000-2020, by both first author publications and last author publications. For both negative binomial models, the outcome variable was the sum of manuscripts published by each race/racism and gender category by publication year. Indicator variables were then constructed for time and to identify Black* authors and women authors. More specifically, the reference categories were men and all other races, centering Black women. When examining the statistical model results, the “Black*” and “women” labels correspond to the average difference as compared to the reference groups.

### MeSH Analysis

To evaluate first authors’ article topics, we paired major MeSH with the first author publications. Percentile rankings for each racial category and associated “feminization” (i.e., higher proportion of women) were calculated using the joint probability for race/racism and the proportion of women authors for individual terms, respectively.

## Results

The sample consisted of 21,945 unique authors. Of these authors, 43% (n = 9,553) were represented as first authors and 37% (n = 8,280) as last. For first authors, 5.6% (n = 541) had missing race/racism values imputed. For last authors, 6.6% (n = 550) had missing race/racism values imputed. Just under five percent (n = 1,090) of authors could not be assigned a gender as their given name was absent from the SSA data.

The first step towards our intersectional analysis was examining the descriptives for race/racism and gender separately from papers published between 2000 and 2020 (see Table 1).

**Table 1.** Summary Statistics on Unique Authors (n = 21,945)

| Category | Statistic |
| --- | --- |
| Black*, Mean (SD) | 0.090 (0.126) |
| Asian*, Mean (SD) | 0.132 (0.276) |
| Hispanic*, Mean (SD) | 0.061 (0.156) |
| Other,* Mean (SD) | 0.024 (0.021) |
| White, Mean (SD) | 0.693 (0.299) |
| Women, % (n)* | 44.9% (9,372) |

\*Gender could not be assigned to 1,090 authors, so the total sample for this category is 20,855.

Black individuals made up only 9% of the sample and women represented 49.9% (excluding non-attributable sources).

Moving to an intersectional lens, Figure 1 presents the extrapolated manuscript counts by author type. Of note, the y-axis has a “floating” (not fixed) scale, which could mislead the reader into thinking that the magnitude of publications across races/racisms are similar. It is not: white women and men have far more publications, followed by Asian* women and men, then Black* women and Black* men, and then Hispanic* women and men. We used this floating scale to highlight the trends for Black* women compared to others and to avoid “drowning out” by white authors. As Figure 2 shows, Black* women have far fewer first and last authorship publications than white women and men. Looking within races, regarding first authorship, Black* women have had higher counts than Black* men since 2015. Similarly, Hispanic* women and Asian* women first authors have larger counts than Hispanic* men and Asian* men respectively.

**Figure 1.**
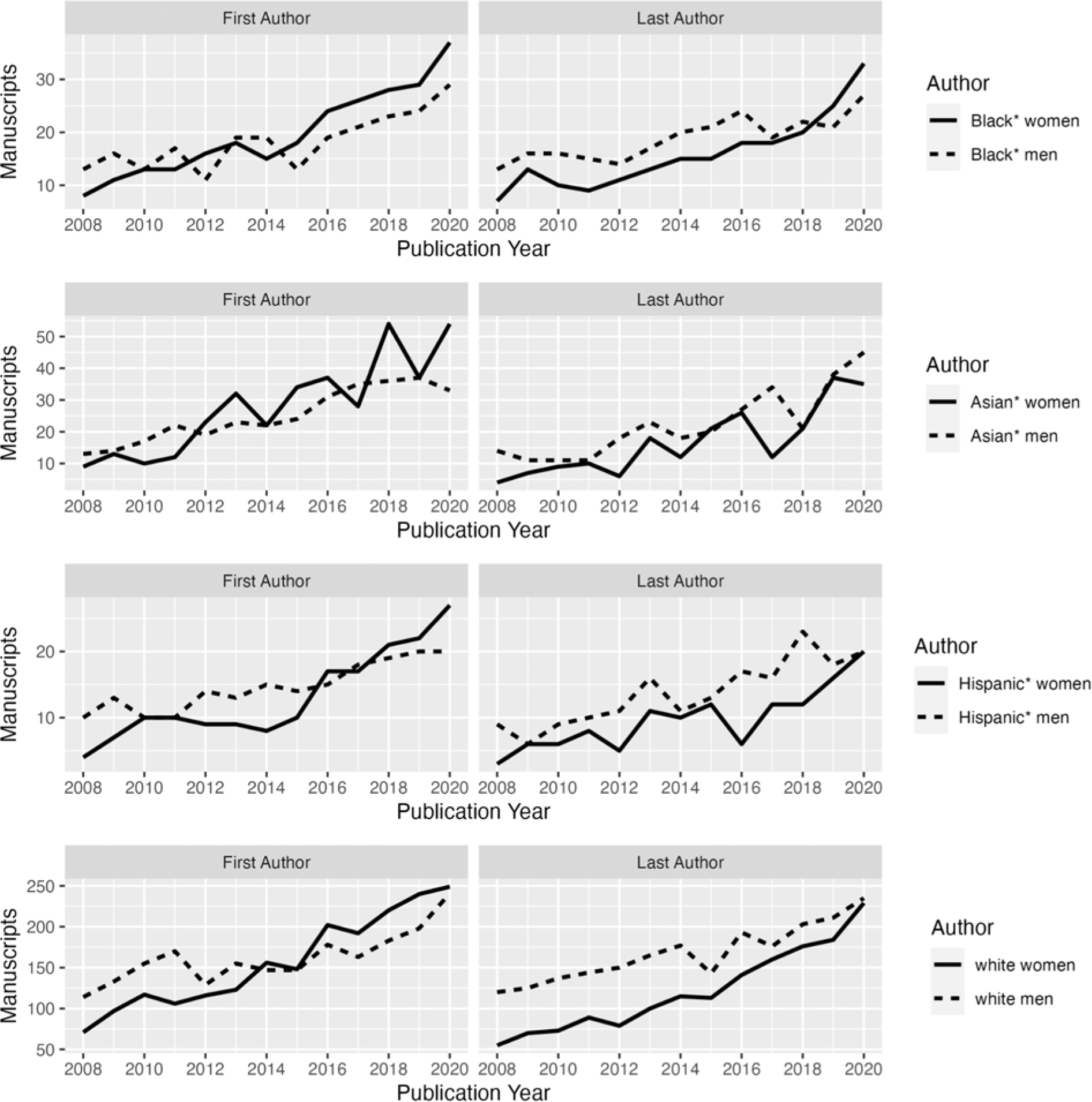
Extrapolated Manuscripts by Author Type, Gender, and Race/Racism. Note: The y-axes in the faceted plots have free-floating scales.

Regarding last authorship, Black* women moved ahead of Black* men in 2018 and Hispanic* women, Asian* women, and white women last authors remain behind Hispanic* men, Asian* men, and white men.

The negative binomial generalized linear model in Table 2, however, shows that the difference between Black* women and Black* men is not statistically significant in either the first or last author model. When examining Black* women and men together, however, there is a significant and large difference: the average incident rate ratio (IRR) among first and last Black* women and men authors, as compared to all other authors, is 66% lower and 64% lower, respectively. In other words, the average publication rate across time decreased by approximately 65% for Black* women and men authors compared to white authors. In both models, the function of time was statistically significant, indicating that the average rate of published manuscripts increases over time.

**Table 2.** Negative Binomial Author Models Results.

| Coefficient | First Author Model |  |  | Last Author Model |  |  |
| --- | --- | --- | --- | --- | --- | --- |
|  | IRR | SE | z Value | IRR | SE | z Value |
| (Intercept) | 29.90 *** | 0.22 | 14.80 | 28.30 *** | 0.23 | 14.2 |
| Black* X<br>Woman | 1.11 | 0.47 | 0.21 | 1.16 | 0.48 | 0.30 |
| Black* | 0.34 ** | 0.33 | -0.31 | 0.36 ** | 0.34 | -2.93 |
| Woman | 0.92 | 0.20 | -0.37 | 0.68 | 0.21 | -1.76 |
| Time | 1.08 ** | 0.02 | 3.08 | 1.09 ** | 0.02 | 3.27 |

\*\*\* = p-value < 0.001; \*\* = p-value < 0.01; \* = p-value < 0.05. IRR = incident rate ratio; SE = standard error.

Regarding topics, the top 10% of major MeSH terms for first authors with greater than 60% feminization and exceeding the 50^th^ percentile ranking in Black* authors (i.e., highly likely Black* women) were: career mobility, cultural diversity, cooperative behavior, manikins, writing, fellowships and scholarships, empathy, internet, and job satisfaction. Comparing these to top major MeSH terms of all authors (Table 3), there appears to be little overlap.

**Table 3.**
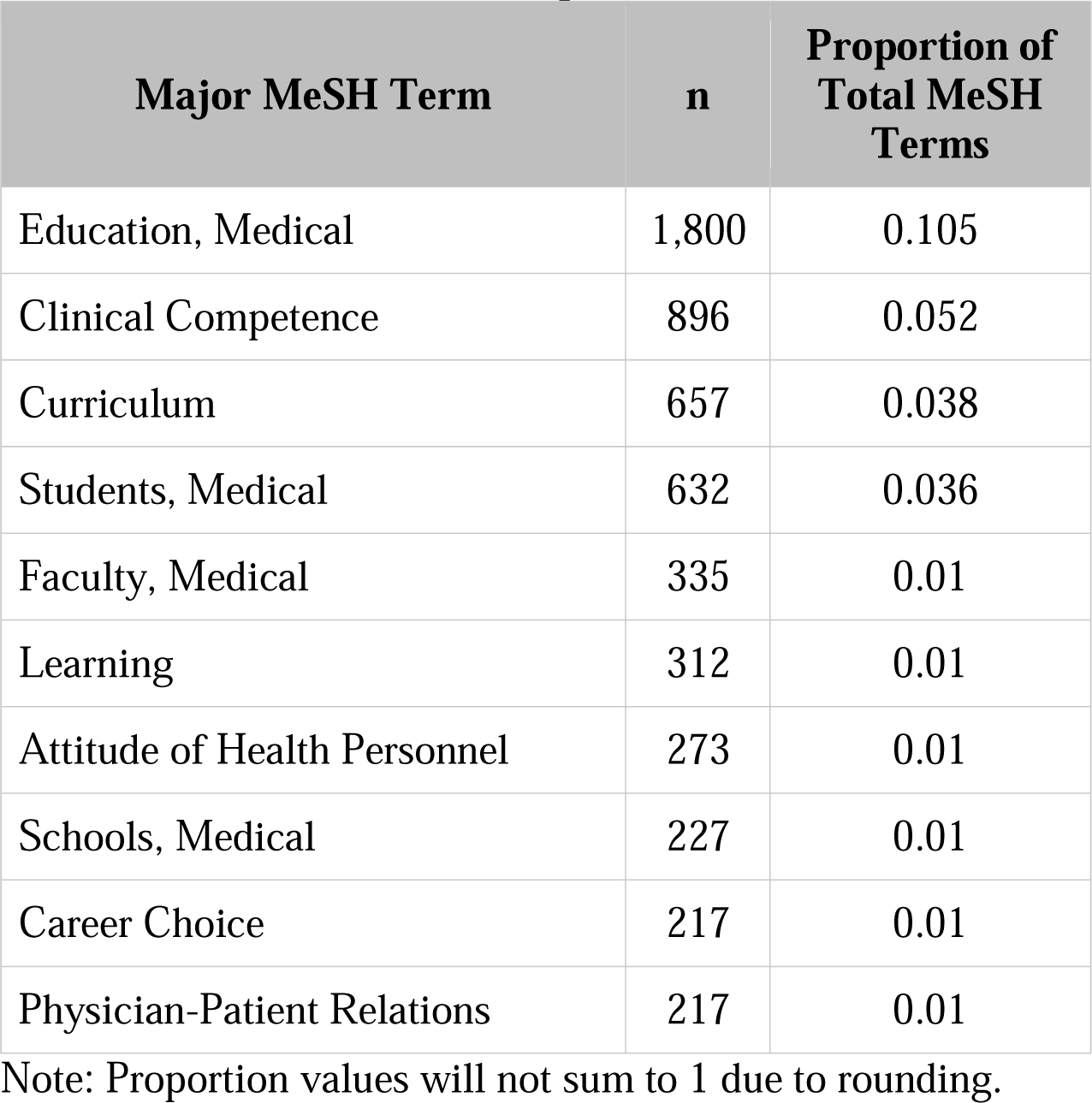
Top 10 Ranking of Overall Medical Subject Heading (MeSH) Terms in Analytic Sample.

First Author **Testimonio** (to contextualize the above statistical account, the first author shares her lived experiences):

### Background: A Little Piece of Haiti

Although born and raised in Queens, NY, my crowded childhood home was a little piece of Haiti. The sounds of Haitian Creole and smells of Haitian food were steeped into my identity as a Black woman. My parents valued education, working multiple jobs around the clock. They expected us to get straight A’s and take up a “respectable” profession: lawyer, doctor, engineer or nurse.

Mirroring my parents’ hard work, I vowed to become a doctor to lead myself out of hardship. This dream seemed far-fetched but my family community supported me, even calling me doctor from the age of 10. This strengthened my resolve despite school counselors recommending that I adjust my goals to be more “realistic.” I went to a vocational high school, commuting one hour each way to get certification as a licensed practical nurse and went on to get a full-ride scholarship to university—my ticket out of Queens. Although I longed to return and practice medicine there, money was a barrier, so I pursued a military medicine scholarship. After completing my Army service obligation as a Pediatrician, I transferred to the Public Health Service looking to have a greater impact on underserved communities. My family are so proud, but still hesitant to accept my westernized medical advice before trying their Haitian remedies. Despite these accomplishments, I feel guilt as one of the few to “make it out.”

### Training and Early Career: Struggling to Belong

The path to medicine was fuzzy to me, but I knew science was integral, so I struggled through hours of bench research, eventually turning in my pipette for a stethoscope and never looked back.

Residency was hard. Despite being intern of the year, I wasn’t sure that I belonged. The nights were long and the work was hard but I was determined to succeed. I tirelessly sought feedback and worked on regaining confidence despite what felt like setbacks.

Upon graduating residency I headed to Germany with my family, instead of doing a chief year or fellowship as previously planned. I made the most of the cards I was dealt and excelled as a general pediatrician, eventually being selected to serve as Chief of Inpatient Pediatrics.

Throughout, teaching medics and others helped me realize I had something to give to academic medicine.

### Academic Medicine: Questioning my Voice’s Worth

When the job at Uniformed Services University came up, it seemed made for me: I would be teaching at the heart of military medicine. Opportunities to teach and mentor were always available. But as I learned more about academic medicine and my role as faculty, I realized my portfolio had a huge gap: research. I reached out to friends and colleagues to discuss ideas for publication in medical education. But nothing materialized, leaving me to believe my ideas weren’t valid even though I would see these same ideas emerge in journals. These experiences and others left me unsure of the worth of my voice in academia.

### Mentoring and Being Mentored: I am More than Enough

Mentoring and empowering medical students to address their concerns regarding their experiences led me to re-establishing myself as an author, addressing issues regarding racism in medicine. Through personal and professional growth, I have realized that I am more than enough and belong in all the spaces I care to be in. Yet I still struggle to be my full self. “Black woman” is a broad label that masks countless ethnicities and identities. My pride and strength in being a Haitian American is masked by this label and all the racism attached to it.

## Discussion

Our IM analysis using QuantCrit tools sought to create a statistical account of Black* women in medical education publishing to supplement experiential accounts. To begin, Black* women are eclipsed by white women and men. This supports Johnson’s and Karvonen et al.’s accounts about their publishing experiences.^4,5^ Further, Author 1’s testimonio demonstrates the experiential effects of this statistical account, emphasizing her struggle for belonging in academic medicine. The exclusion of Black* women’s voices and the experiential effects of that exclusion denies them equal access to the resources to create knowledge. This is something any leader in medical education publishing must claim responsibility for. Authors 2, 3, and 5 are editors contributing to this exclusion. Following Haynes’ focus on institutional change, we commit to partnering with Black* women within our own editorial work through actions such as tracking Black* women’s publication trends, actively soliciting manuscripts from Black women authors, or intentionally amplifying and sponsoring Black* women in our field to publish. We call for others to do the same.

Furthermore, this study found that none of the top 10 MeSH topics common to Black* women authors appeared within the top 10 overall ranking of topics in medical education. In fact, the two topics Black* women authors most addressed were career mobility (defined as “the upward or downward mobility in an occupation or the change from one occupation to another”^42^) and cultural diversity (described as the “coexistence of numerous distinct ethnic, racial, religious, or cultural groups within one social unit, organization, or population”--notably, a parent term for diversity, equity, inclusion^43^). Author 1’s testimonio further contextualizes this: her entry into medical education publishing was on the topic of diversity. The focus on these topics raises questions about potential gendered/racialized topic expectations, resulting in an additional minority tax levied upon Black* women authors as ‘Superwomen’.^1,11^ Researchers’ topic selection behaviors are influenced by multiple intersecting factors, including gender, race/racism, research interests and background, and funding opportunities.^45^ However, research topic selection comes at a cost, with topics covered by Black* women authors garnering fewer citations compared to other groups.^14^ These findings have critical implications for Black* women researchers’ career success and trajectory and harms the field without the full and unfettered participation of Black* women.

This study must be considered in light of its limitations. First, our methodology essentialized and simplified gender and race/racism, so we could not fully instantiate IM. Racial and gender identification is manifold and dynamic (e.g., two or more races/racisms; nonbinary or shifting gender identity); applying an aggregate-level algorithm likely resulted in misattribution of racial and gender categories. Second, data were collected for only researchers identifying as based at US institutions, posing interpretation issues for global authors in the sample. Additionally, some authors may not have been born in the US, impacting SSA data, which resulted in over 1,000 missing name values. Third, we used the MEJ-24, which does not include medical education articles published in journals not specific to medical education (e.g., clinical journals). Moreover, we were reliant on what data publishers—largely *not* Black* women— chose to collect and how they collected it.

This study suggests that the harm to Black* women in the broader field of medical education^3^ holds in publishing as well. Authorship–the “coin of the realm” in academia^45^–is a central piece of this puzzle. Quantitative accounts like this one help us to understand the scope of the harm. Yet we must also draw other Black* women—those who already are and desire to be authors—into dialogue through experiential accounts. How do they experience authorship in medical education and how can we ensure that they do not feel, as Johnson did amidst peer review, that “’*you* are not enough,’ ‘your *story* is not enough,’ ‘your *voice* is not enough.’”?^4^

## Acknowledgements

We acknowledge and thank Joseph Costello for his assistance in formatting this manuscript.

## Disclaimer

The views expressed in this article are those of the authors and do not necessarily reflect the official policy or position of the Uniformed Services University of the Health Sciences, the Department of Health and Human Services, the Department of Defense, or the U.S. Government.

## References

1. Balzora S. When the minority tax is doubled: being Black and female in academic medicine. Nat Rev Gastroenterol Hepatol. 2021;18(1):1.

2. Rezaiefar P, Abou-Hamde Y, Naz F, Alborhamy YS, LaDonna KA. “Walking on eggshells”: experiences of underrepresented women in medical training. Perspect Med Educ. 2022;11(6):325–332.

3. Association of American Medical Colleges. Diversity in Medicine: Facts and Figures 2019. https://www.aamc.org/data-reports/workforce/report/diversity-medicine-facts-and-figures-2019. Accessed August 30, 2023.

4. Johnson M, Konopasky A. Maintaining your voice as an underrepresented minority during the peer review process: A dialogue between author and mentor. Perspect Med Educ. 2022;11(3):144–145.

5. Karvonen KL, Bonachea EM, Burris HH, et al. Addressing bias and knowledge gaps regarding race and ethnicity in neonatology manuscript review. J Perinatol. 2022;42(11):1546–1549.

6. Rice DB, Raffoul H, Ioannidis JPA, Moher D. Academic criteria for promotion and tenure in biomedical sciences faculties: cross sectional analysis of international sample of universities. BMJ. 2020;369:m2081.

7. Covarrubias A, Nava PE, Lara A, Burciaga R, Vélez VN, Solorzano DG. Critical race quantitative intersections: a testimonio analysis, Race Ethnicity and Education. 2018;21(2):253–273. doi:10.1080/13613324.2017.1377412

8. Madden C, O’Malley R, O’Connor P, O’Dowd E, Byrne D, Lydon S. Gender in authorship and editorship in medical education journals: A bibliometric review. Med Educ. 2021;55(6):678-688.

9. Maggio LA, Costello JA, Ninkov AB, Frank JR, Artino AR Jr. The voices of medical education scholarship: Describing the published landscape. Med Educ. 2023;57(3):280–289.

10. Gillborn D, Warmington P, Demack S. QuantCrit: Education, policy,‘Big Data’and principles for a critical race theory of statistics. Race Ethnicity and Education. 2018;21(2):158–79. doi:10.1080/13613324.2017.1377417

11. Bajaj SS, Tu L, Stanford FC. Superhuman, but never enough: Black women in medicine. Lancet. 2021;398(10309):1398–1399.

12. Ajjawi R, Crampton PES, Ginsburg S, et al. Promoting inclusivity in health professions education publishing. Med Educ. 2022;56(3):252-256.

13. Anna T. Letter from the Editor - Teaching & Learning in Medicine’s Anti-Racism Strategy. Teach Learn Med. 2020;32(5):457-458.

14. Kozlowski D, Larivière V, Sugimoto CR, Monroe-White T. Intersectional inequalities in science. Proc Natl Acad Sci USA. 2022;119(2):e2113067119.

15. Mohammadi E, Gregory KB, Thelwall M, Barahmand N. Which health and biomedical topics generate the most Facebook interest and the strongest citation relationships? Information Processing & Management. 2020;57(3):102230. doi:10.1016/j.ipm.2020.102230

16. Hopkins AL, Jawitz JW, McCarty C, Goldman A, Basu NB. Disparities in publication patterns by gender, race and ethnicity based on a survey of a random sample of authors. Scientometrics. 2013;96:515–34. doi:10.1007/s11192-012-0893-4

17. Nunkoo R, Thelwall M, Ladsawut J, Goolaup S. Three decades of tourism scholarship: Gender, collaboration and research methods. Tourism Management. 2020;78:104056. doi:10.1016/j.tourman.2019.104056

18. Plowman DA, Smith AD. The gendering of organizational research methods: Evidence of gender patterns in qualitative research. Qualitative Research in Organizations and Management. 2011;6(1):64–82. doi:10.1108/17465641111129399

19. Thelwall M, Bailey C, Tobin C, Bradshaw NA. Gender differences in research areas, methods and topics: Can people and thing orientations explain the results? Journal of Informetrics. 2019;13(1):149–169. doi:10.1016/j.joi.2018.12.002

20. Thelwall M, Abdoli M, Lebiedziewicz A, Bailey C. Gender disparities in UK research publishing: Differences between fields, methods and topics. Information Professional. 2020;29(4):e290415. 10.3145/epi.2020.jul.15

21. Crenshaw K. Demarginalizing the Intersection of Race and Sex: A Black Feminist Critique of Antidiscrimination Doctrine, Feminist Theory and Antiracist Politics. University of Chicago Legal Forum. 1989;1989(1):8. Available at: http://chicagounbound.uchicago.edu/uclf/vol1989/iss1/8

22. Zuberi T. Deracializing social statistics: Problems in the quantification of race. In: Zuberi T, Bonilla-Silva E, White Logic, White Methods: Racism and Methodology. Maryland: Rowman & Littlefield Publishers; 2008:127-134.

23. Garcia NM, López N, Vélez VN. QuantCrit: Rectifying quantitative methods through critical race theory. Race Ethnicity and education. 2018;21(2):149–57. doi:10.1080/13613324.2017.1377675

24. Haynes C, Joseph NM, Patton LD, Stewart S, Allen EL. Toward an understanding of intersectionality methodology: A 30-year literature synthesis of Black women’s experiences in higher education. Review of Educational Research. 2020;90(6):751–87. doi:10.3102/0034654320946822

25. Zuberi T, Bonilla-Silva E. White Logic, White Methods: Racism and Methodology. Lanham, MD;Rowman & Littlefield Publishers Inc.; 2008.

26. Zaidi Z, Rockich-Winston N, Chow C, Martin PC, Onumah C, Wyatt T. Whiteness theory and the (in)visible hierarchy in medical education. Med Educ. 2023;57(10):903–909.

27. Berry C, Khabele D, Johnson-Mann C, et al. A Call to Action: Black/African American Women Surgeon Scientists, Where are They?. Ann Surg. 2020;272(1):24–29.

28. Ighodaro ET, Littlejohn EL, Akhetuamhen AI, Benson R. Giving voice to Black women in science and medicine. Nat Med. 2021;27(8):1316–1317.

29. Yip SWL, Rashid MA. Editorial diversity in medical education journals [published correction appears in Clin Teach. 2022;19(2):180]. Clin Teach. 2021;18(5):523-528. doi:10.1111/tct.13386

30. Tsai J, Lindo E, Bridges K. Seeing the Window, Finding the Spider: Applying Critical Race Theory to Medical Education to Make Up Where Biomedical Models and Social Determinants of Health Curricula Fall Short. Front Public Health. 2021;9:653643.

31. Zaidi Z, Partman IM, Whitehead CR, Kuper A, Wyatt TR. Contending with Our Racial Past in Medical Education: A Foucauldian Perspective. Teach Learn Med. 2021;33(4):453–462.

32. Squazzoni F, Bravo G, Farjam M, et al. Peer review and gender bias: A study on 145 scholarly journals. Sci Adv. 2021;7(2):eabd0299.

33. Salazar JW, Claytor JD, Habib AR, Guduguntla V, Redberg RF. Gender, Race, Ethnicity, and Sexual Orientation of Editors at Leading Medical and Scientific Journals: A Cross-sectional Survey. JAMA Intern Med. 2021;181(9):1248-1251.

34. Strmic-Pawl HV, Jackson BA, Garner S. Race counts: racial and ethnic data on the US Census and the implications for tracking inequality. Sociology of Race and Ethnicity. 2018 Jan;4(1):1–3. doi:10.1177/2332649217742869

35. Robinson Jr J. Live Black Retire Poor Die Early: How Social Security as an Institution Continues to Perpetuate the Social Racism of the 1930s. Elder LJ. 2016;24:487.

36. A. Ninkov, Frank JR, Maggio LA. Bibliometrics: Methods for studying academic publishing. Perspect Med Educ. 2022;11(3):173-176.

37. Maggio LA, Ninkov A, Frank JR, Costello JA, Artino AR Jr. Delineating the field of medical education: Bibliometric research approach(es). Med Educ. 2022;56(4):387–394.

38. Zitt M, Lelu A, Cadot M, Cabanac G. Bibliometric Delineation of Scientific Fields. In: Glänzel, W., Moed, H.F., Schmoch, U., Thelwall, M. (eds) Springer Handbook of Science and Technology Indicators. Springer Handbooks. Springer, Cham. doi: 10.1007/978-3-030-02511-3_2

39. Muñoz-Écija T, Vargas-Quesada B, Rodríguez ZC. Coping with methods for delineating emerging fields: Nanoscience and nanotechnology as a case study. Journal of Informetrics. 2019;13(4):100976. doi:10.1016/j.joi.2019.100976.

40. Lietz H. Drawing impossible boundaries: field delineation of Social Network Science. Scientometrics. 2020;125:2841–2876. doi:10.1007/s11192-020-03527-0.

41. U.S. Social Security Administration. Popular Baby Names: Beyond the Top 1000 Names. https://www.ssa.gov/oact/babynames/names.zip. Accessed: December 1, 2022.

42. Career Mobility. Medical Subject Headings (MeSH. United States: National Library of Medicine; 1963. https://www.ncbi.nlm.nih.gov/mesh/68002322. Updated June 8, 2023. Accessed August 30, 2023.

43. Cultural Diversity. American Heritage Dictionary, 2d ed. 1982:p955 Cited by: Cultural Diversity. Medical Subject Headings (MeSH). United States: National Library of Medicine; 1963. https://www.ncbi.nlm.nih.gov/mesh/68018864. Updated June 8, 2023. Accessed August 30, 2023.

44. Zhang C, Wei S, Zhao Y, Tian L. Gender differences in research topic and method selection in library and information science. Library & Information Science Research. 45(3):101255.

45. Wilcox LJ. Authorship: the coin of the realm, the source of complaints. JAMA. 1998;280(3):216–217.

